# A GENOMIC BASIS FOR TRANS-OCEANIC SEA TURTLE MIGRATION

**DOI:** 10.64898/2026.08.18.745595

**Authors:** Jamie Adkins, Abdul Hamid A. Toha, Deasy Lontoh, Fitry Pakiding, Andhika P. Prasetyo, Peter H. Dutton, Ekaterina Osipova, Jeffrey A. Seminoff, Tomoharu Eguchi, Scott R. Benson, Lisa M. Komoroske

## Abstract

Long-distance migration has evolved repeatedly across the animal kingdom, yet the underlying processes giving rise to and maintaining these complex eco-behavioral phenotypes remain poorly understood. Here, we present the first evidence of genomic determinants of migratory phenotypes in sea turtles, using whole genome resequencing to demonstrate that complex genomic architecture underlies divergent migratory destinations and reproductive timing in the critically endangered western Pacific leatherback turtle (*Dermochelys coriacea*). Individuals from this admixed population that navigate to foraging grounds on opposite sides of the Pacific Ocean have a putative inversion on chromosome 2 encompassing one gene, potentially conferring pleiotropic physiological effects and supporting magnetoreception. Genomic architecture underlying divergent reproductive timing is more dispersed, aligned with reduced gene flow, and is associated with genes that may influence reproductive success. Genes underlying both traits suggest a role for neurodevelopment and memory. Our study adds to the increasing evidence of at least partial genomic control of migratory traits in wild populations, with important potential implications for conservation measures such as translocation and genetic rescue. Our results align with a growing body of work describing complex genomic architecture and structural variants underlying key eco-behavioral traits, advancing the understanding of evolution of long-distance migration across taxa.

## INTRODUCTION

Long-distance migration is a critical component of life history for many animal taxa, enabling organisms to overcome spatiotemporal variation in resources and environmental conditions to shape their evolution and ecology [1,2]. Migratory phenotypes are governed by complex physiological, genetic, and environmental interactions, and can exhibit variation between and within populations for traits such as orientation, timing, and propensity to migrate [3]. To successfully migrate, animals must have a spatiotemporal migration program, including an internal timing system and known migratory direction [4]. Such a program is likely inherited and further shaped by genome-environmental interactions, such as learning and memory processes modulated by environmental cues for migratory traits such as timing (e.g. temperature, day length) and orientation (e.g. polarized light, geomagnetic field) [1]. While the genetic and environmental drivers of migration have long been topics of interest in evolutionary biology and animal physiology, only recently have technological advances enabled examination of these factors in free-ranging migratory species. Studies have now identified genetic associations underlying migratory phenotypic diversity in fishes, birds and insects, with comparisons across taxa generally finding different molecular signatures of migratory phenotypes between evolutionary lineages [2]. However, this work has only been conducted on a small proportion of migratory species with some lineages completely unstudied, so it remains unclear if molecular mechanisms driving migratory traits are shared or unique across taxonomic groups. Additionally, 44% of all migratory species globally are in decline [5], and their ability to cope with rapid environmental change is likely influenced by the mechanisms driving their migratory behaviors (e.g., fixed or flexible responses to changes in environmental cues or breeding habitat suitability). Therefore, continued work within and across lineages is needed to advance our comparative understanding of the genomic drivers of migratory phenotypes and determine their roles in species risk and resilience.

The rise and maintenance of complex phenotypes, including migratory behaviors, may be governed by various genomic architectures that are still poorly understood in naturally occurring populations. For example, it has been widely assumed that complex eco-behavioral phenotypes must be driven by many loci of small effect distributed across the genome, but recent studies have found evidence of a few loci or supergene regions with large effects in migratory phenotypes [6–9]. Studies of other ecotypes have come to similar conclusions, supporting an alternative hypothesis that complex traits can be controlled by a few loci of large effect, either alone or alongside loci of smaller effect [10]. A related active area of interest is understanding how such divergent phenotypes are preserved in scenarios of high gene flow, which should have a homogenizing effect on genomic variation [11]. Increasingly, studies across diverse taxa have identified genomic islands of divergence linked to ecological adaptation and phenotype differences that arise and are maintained despite little to no differentiation across the rest of the genome [12]. The genetic divergence in these regions may be driven by several mechanisms, such as clustering of loci in low-recombination regions [13], structural variants (e.g., inversions and supergenes) [14], genetic hitchhiking due to strong divergent selection [15], or a combination of these processes[16]. Structural variants are suggested to be particularly important to the maintenance of divergent phenotypes in scenarios of high gene flow [17], with examples including trout [18], *Littorina saxatilis* wave and crab ecotypes [19], and migratory birds [20]. While the roles of structural variants and these processes are still unknown for many eco- behavioral phenotypes, recent advances in high-quality reference genomes and whole genome resequencing are now enabling these studies more broadly in wild populations.

Sea turtles are an iconic migratory taxon, well-known for their oceanic journeys between distant nesting and foraging grounds [21]. Although all sea turtle species exhibit migratory behaviors, the physiological mechanisms and underlying genetic control of these phenotypes remain unknown. Within Sauropsids, sea turtles share a recent common ancestor with birds, but represent a unique study system as marine, non-avian reptiles. Like migratory birds, sea turtles likely utilize a combination of macro-scale (geomagnetic inclination and intensity) and fine-scale (chemical and olfactory) cues to locate distinct natal beaches [22], suggesting that variation in sensory processes and spatial memory exists within and among populations. Geomagnetic “map” and “compass” senses are likely to be at least in part inherited as sea turtles receive no parental care, are primarily solitary, and must navigate novel seascapes with few identifiable landmarks to habitats that change during their long lifespan [23]. While their avian relatives may also utilize geomagnetic cues for navigation, these differences in environmental conditions and life history traits suggest different biological and evolutionary processes may underlie mechanisms of long-distance migration in sea turtles.

Leatherback turtles (*Dermochelys coriacea*) undertake the longest migrations of all sea turtles, repeatedly completing trans-oceanic migrations over their lifetime [24]. In the critically endangered western Pacific leatherback metapopulation [25] , a significant proportion of nesting occurs on a single peninsula in Indonesia with individuals displaying divergent migratory phenotypes in reproductive timing and post-nesting migratory destination [26,27] (Fig. 1). Females undertake nesting re-migrations every 2–6 years [28], returning to their natal beaches in the boreal winter (wet season, hereafter “winter”) or boreal summer (dry season, hereafter “summer”) months. Satellite telemetry and stable isotope analyses of post-breeding individuals indicate that winter nesters migrate to foraging grounds primarily in the southern hemisphere, including the East Australian Current (EAC) and Tasman Sea, or nearby Indonesian seas, whereas summer nesters largely utilize pelagic foraging grounds in the northern hemisphere, mainly the proximate South China Sea (SCS) or distant California Current Ecosystem (CCE) [26,27] and possibly the North Pacific Transition Zone-Kuroshio Extension [26,28]. Therefore, winter and summer nesters do not co-occur at nesting beaches or foraging grounds, and individuals exhibit fidelity to foraging sites, as in other sea turtle species ([29]; S. Benson, pers comm. 2026). While these behaviors are undoubtedly influenced by environmental factors [30], the ability to find productive foraging areas in large and highly heterogeneous oceanic habitats is likely to be evolutionarily favored and driven by genomic variation [31] and/or genomic- environmental interactions (e.g., the learned migration goal hypothesis [30]). Though the origins or molecular drivers of these divergent migratory behaviors are unknown, different environmental conditions between the regions may contribute to maintaining them. For migratory destination, the temperate CCE has dense seasonal jellyfish blooms [26], their primary food, potentially conferring a growth and reproductive advantage that overcomes the greater energetic costs associated with trans-oceanic travel. For reproductive timing, seasonal reversals in the direction of the New Guinea Coastal Current may affect hatchling dispersal [30], contributing to isolation of individuals hatched in summer versus winter months into northern and southern hemisphere habitats, respectively, facilitating demographic independence via temporal reproductive isolation (Fig. 1). However, these current reversals do not always occur [30], and studies to date have not found evidence of genetic structure in this population [27,32]. Thus, while environmental factors likely play important roles in migratory and reproductive behaviors, we hypothesize that there are inherited genetic components to the divergent migratory destination and reproductive timing phenotypes observed in leatherback turtles.

**Figure 1.**
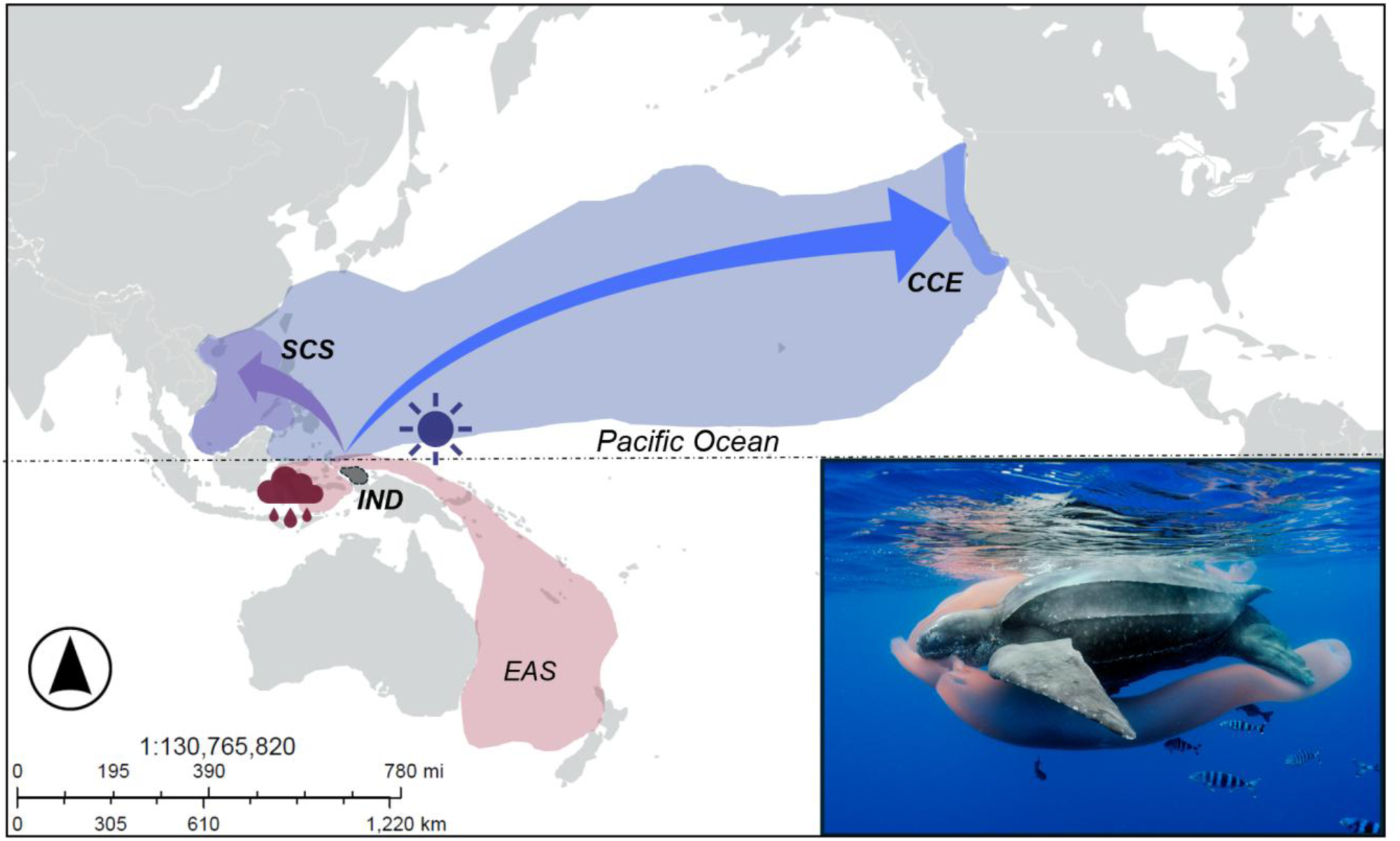
Map of western Pacific leatherback turtle nesting and foraging habitat and migratory connections. Grey shaded area indicates the primary nesting beaches in Indonesia (IND) for this population. Boreal summer nesters utilize foraging areas in the northern hemisphere (blue shaded region north of dashed equator line), including the South China Sea (SCS, purple shading) and California Current Ecosystem (CCE, blue shading). Boreal winter nesters migrate to southern hemisphere foraging grounds (designated by the red shaded area south of the dashed equator line), including the East Australian Current (EAC). Image: leatherback turtle foraging on gelatinous prey; courtesy of Brian Skerry.

Here, we provide the first study of sea turtle migratory genomics using western Pacific leatherbacks as a study system. Given the high gene flow within this population and distinct divergent phenotypes, we predicted that a genomic basis governing migratory phenotypes would likely involve complex genomic architecture. Combining whole genome resequencing of individuals with known migratory phenotypes via satellite telemetry and stable isotope assignment [26,33], we (1) characterize the genomic basis of leatherback migratory destination and reproductive timing; (2) determine how these complex migratory phenotypes would be maintained under a high gene flow scenario; and (3) identify biological processes putatively associated with long-distance migration using a chromosomal level reference genome with species-specific annotation [34]. Our study provides the first evidence of at least partial genomic control of migratory phenotypes in sea turtles with implications for their responses to environmental change and other anthropogenic pressures, and also more broadly contributes to our understanding of the evolution and maintenance of complex eco-behavioral phenotypes in natural systems.

## METHODS

### (a) Study area and design

We analyzed whole genome resequencing data (WGR) [27] of adult female western Pacific leatherback turtles nesting at Bird’s Head Seascape, Papua-Indonesia sampled across 9 years (2003-2011) with corresponding individual migratory phenotypic information. Available nesting sites during the summer include the larger Jeen Yessa (∼17.9km) and smaller Jeen Syuab (∼ 6km) beaches, which are separated by cliffs and rocky outcroppings; seasonal monsoons erode the entire beach at Jeen Yessa, restricting winter nesting to Jeen Syuab beach. For migratory destination comparisons, genetic samples paired satellite telemetry or stable isotope assignment to foraging ground was only available for summer nesters (n=48; [26]); of these we included individuals that clearly assigned to either the SCS (n=21) or CCE (n=25) foraging grounds. For reproductive timing comparisons, winter nesters (n=12) were compared to a subset of summer nesters from all the individuals used in the migratory destination analysis, randomly selected within each foraging ground (SCS: n=6; CCE: n=6). To independently validate associations between eastern migratory destination phenotype and genotypes, we performed WGR of 10 individuals sampled while foraging in the CCE across 19 years (2003-2022). Validation samples were collected during routine in-water population monitoring by the NOAA Southwest Fisheries Science Center (approved activities under NOAA-ESA permit numbers 1227, 15634, and 21111) as described in [35]. WGR libraries were prepared using an Illumina Nextera PCR-based protocol adjusted for one tenth reaction sizes, as described in [35], and sequenced on a NovaSeqX lane with 150PE chemistry at Novogene Corporation, Inc (Sacramento, CA, USA).

### (b) Bioinformatic processing

All code used for analyses is available as described in the data accessibility section. Briefly, we assessed sequence data quality (Fastqc v0.11.9 [36]), trimmed sequences (bbduk v38.90 [37]), and aligned reads (BWA-mem v0.7.17 [38]) to the *Dermochelys coriacea* reference genome (GCA_009764565.4 [34), sorted and indexed files (Samtools v1.14 [39]), marked duplicates (Picard v2.27.2 [40]), and assessed depth of coverage (Mosdepth v2.26.2 [41]). Samples with mean coverage <1X were removed from further analysis. For all samples with mean coverage >6X, we also used GATK v4.2.3.0 to call SNPs following GATK best practices for workflow and filtering [42,43].

### (c) Identification of Genomic Islands of Divergence

Genomic islands of divergence (GID) are highly differentiated contiguous regions containing linked SNPs in a region of the genome that otherwise exhibits low differentiation [44]. We first implemented a genome-wide sliding window F_st_ scan using genotype likelihoods (ANGSD v0.935; Korneliussen et al. 2014) including all samples that passed initial quality filter thresholds. For each phenotypic comparison (migratory destination n=48; reproductive timing n=24) we generated a list of all SNPs passing filters for quality, minimum minor allele frequency (MAF), individual and depth. Then for each phenotype group, we generated site allele frequency (SAF) and MAF files restricted to SNPs present in the site list, with filters as above but without MAF or snpPval filters as inclusion of these filters can impact inferences of genotype-phenotype associations [45]. We computed the folded site frequency spectrum (SFS), and used this to compute F_st_ values in sliding windows of 10kb across the genome. We defined F_st_ outliers as those within the 99.9^th^ percentile of all windows. To better understand the evolutionary processes potentially driving patterns of relative genomic differentiation between phenotypes, we also computed absolute nucleotide divergence (D_xy_) for all SNPs using ngsPopGen [46] in R v4.3.0 [47].

### (d) Genotype-phenotype associations

Since F_st_ peaks in the genome can be the result of multiple evolutionary processes, we also identified outlier SNPs associated with migratory phenotypes using statistics that allow for binomial comparison of SNPs between groups (Baypass v2.2 [48]). We incorporated genotype likelihoods by converting the MAF at each site to allele count data format, and then used a standard covariate model to generate X^t^X and C_2_ statistics as well as Bayes factor values for each SNP using a binary contrast to define phenotype (i.e., foraging ground assignment or nesting season). We ran three iterations of baypass with random seed values as recommended for MCMC methods. We took the median value of each statistic to identify SNP associations to phenotypes, and calibrated each statistic using pseudo-observed datasets [48]. We finally generated a final conservative list of outlier SNPs for each phenotype compassion by including only those that were within the 99^th^ percentile across X^t^X, C_2_ , and F_st_ analyses.

### (e) Functional analysis

To characterize the functional genomic basis of migratory direction and reproductive timing phenotypes, we extracted annotation information (e.g., intron, exon) for outlier SNPs from the reference genome (SNPeff v2017-11-24 [49]), supplemented with functional data from public databases and literature searches for outlier genes. To identify the biological processes associated with outlier genes we used the Panther database (www.pantherdb.org; accessed Jan 2024), and performed enrichment analysis (Metascape v3.5.20240901 [50]), with genes mapped to *H. sapiens* gene IDs as the gene universe to retain as much functional information as possible.

### (f) Structural Variant identification

Emerging work indicates differences in genomic architecture can be important to maintain ecotypes in populations with high levels of gene flow [14]. To determine if structural variants contribute to maintenance of leatherback divergent ecotypes, we performed local sliding window PCA analyses (R package lostruct v0.0.0.9 [51]). Outlier regions were chosen for local PCA if at least 5 outlier SNPs were in a continuous region on a chromosome. Regions were extended by 1Mb before and after a given outlier region to aid in identifying inversion breakpoints. For individuals with mean coverage >6X, we extracted hard-called genotypes for all SNPs within these extended regions using VCFtools (v0.1.16; [42], which were used as input for conversion to genofile format with (R package SNPrelate v1.36.1 [52]). We performed sliding window PCA and multidimensional scaling (MDS) to identify the location of putative inversions, and mapped outlier SNPs to putative inverted regions to assess degree of overlap. To identify regions of elevated linkage disequilibrium (LD) expected with inversions, we computed the r^2^ statistic in windows thinned by 1000 (VCFtools v0.1.16; [42]), and visualized values along the chromosomal region of interest (R package ldheatmap v1.0-5 [53]). Finally, we computed heterozygosity per individual across the putative inverted region, ran a single PCA on the putative inverted region and plotted this against heterozygosity to see if individuals clustered into three distinct groups corresponding to inversion genotypes. We defined regions as putative inversions if they met the criteria of: (1) having distinct peaks when visualizing local PCA and MDS for the region, (2) elevated LD with clear boundaries, and (3) three clear clusters when visualizing the first principal component of local PCA against heterozygosity for the region. Putative inversions were categorized as small: <100Kb, intermediate: 100-500Kb, large: >500Kb [54]).

## RESULTS

### (a) Oligogenic genomic architecture of migratory destination

After filtering and variant calling, we retained a set of 3.2 million SNPs and 58 individuals with a median coverage=15.92x (Table S1) for downstream analyses. We found consistent, strong support for genomic islands of divergence associated with migratory destination (i.e., SCS vs. CCE foragers; Fig. 2A; Table S2). We identified clear F_st_ peaks spanning regions up to 1Mb in length, differentiating individuals foraging in the SCS or CCE, with highest F_st_ on chromosomes 2, 4, 10, and 19 (F_st_ > 0.2; Table S2). We found 957 and 165 outlier SNPs for C_2_ and X^T^X tests, respectively (Table S2). Our final outlier list resulted in 111 SNPs distributed across 13 chromosomes. However, these outlier SNPs were not uniformly distributed across chromosomes – 76 (68.4%; Table S2) occurred within one gene on chromosome 2 (Figs. 2A, S1), indicating a GID and that the genomic architecture of this trait is oligogenic rather than polygenic. D_xy_ was high (99.9^th^ percentile; Table S2, Fig. S1) for 78% of outlier SNPs (median = 0.58; 0-1 scale).

**Figure 2.**
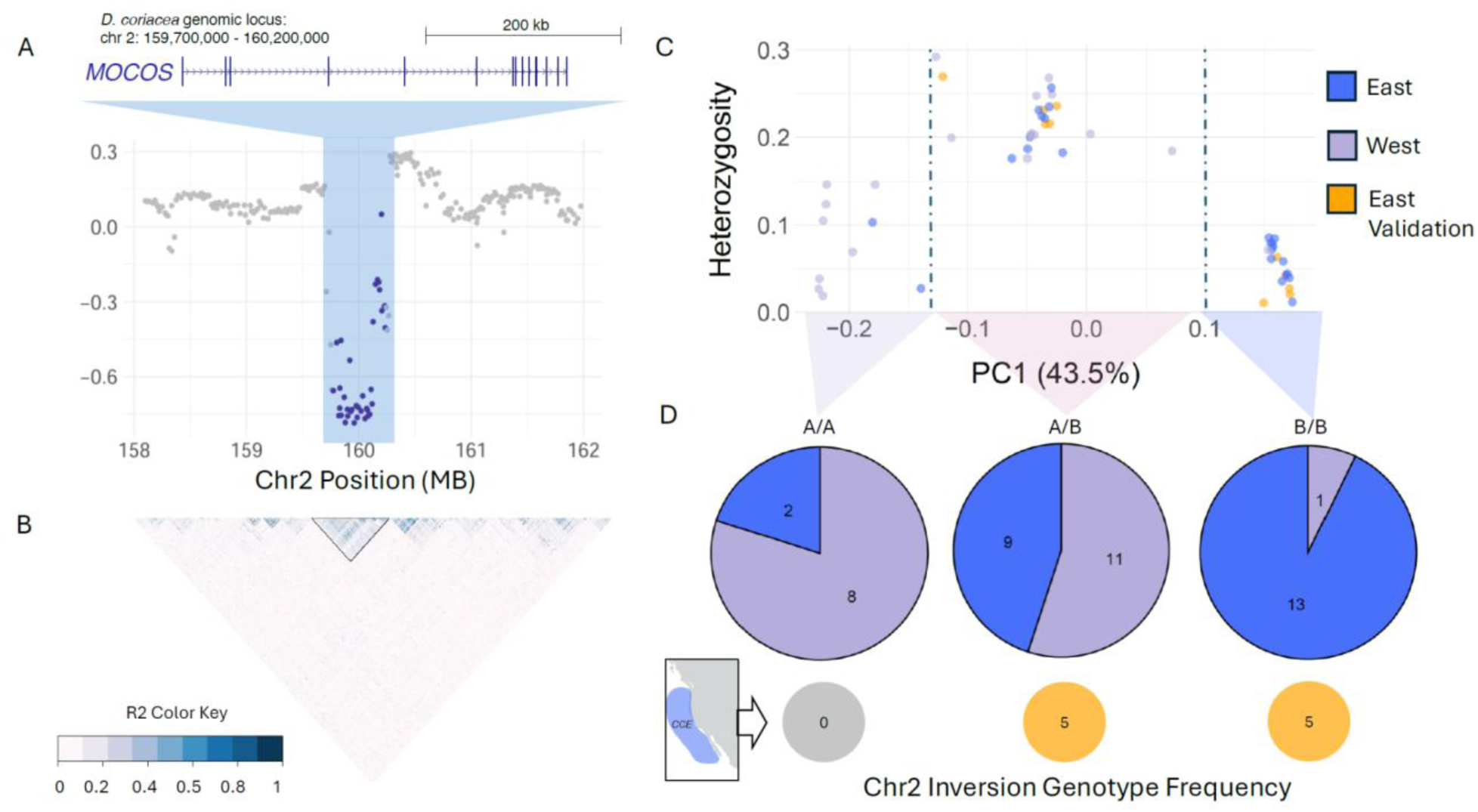
For migratory destination phenotype A) MDS plot of the putative chromosome 2 inversion, X axis is genome position in Mb. The *Mocos* gene contained within the inversion is indicated at the top. Purple points indicate 50bp windows where outlier SNPs occur, blue box denotes putative inversion region. B) LD measured as r^2^ across a portion of chromosome 2 that encompasses the putative inversion. The black triangle outline and indicates a high LD block at the inverted region. Dark colors indicate elevated LD, light purple indicates low LD. C) Plot of the first principal component (PC1) from PCA of SNPs within the putative chromosome 2 inversion (X-axis) versus heterozygosity in that region (Y-axis). Individuals grouped into three clusters (separated by the dashed lines), with the left cluster representing one homozygous inversion genotype, the middle cluster with high heterozygosity indicating individuals with heterozygous inversion genotype, and the right-most cluster indicating the alternate homozygous inversion genotype.

Further examination of allele frequencies at the 111 outlier SNPs suggested that no single SNP is driving divergence between phenotypes (Table S2, Figs. S2 & S3).

### (b) Structural variation underlies divergent trans-oceanic migratory destinations

We identified 4 regions of contiguous outlier SNPs on chromosomes 2, 4, 10, and 19 (Table S2). Of these, we classified the chromosome 2 region as a putative inversion of intermediate size spanning 470Kb, or 0.1% of chromosome 2 (approximately bp 159713092-160279450; Fig. 2A). Local PCA of this region explained 30.5% (PC1) and 12.7% (PC2) of variance (Fig. S3). This region displayed expected signatures of local recombination suppression characteristic of inversions [55], including a continuous, elevated linkage disequilibrium (LD) along the length of the inverted region with clear boundaries (Fig. 2B), and PCA clustering of individuals into three distinct groups with elevated heterozygosity estimates for heterozygous individuals (Fig. 2C). In contrast, outlier regions on chromosomes 4, 10, and 19 did not meet all putative inversion criteria. Together, these results strongly support an inversion on chromosome 2, but not on other chromosomes, underlying this phenotype.

### (c) Inversion genotypes were associated with foraging phenotype

Homozygous individuals for the chromosome 2 inversion clustered strongly by foraging ground. For simplicity, we denote the two inversion types as “Type A” and “Type B”. Of individuals with the homozygous A/A inversion genotype, 80% had west (SCS) foraging phenotypes; similarly, 92.8% of the individuals with B/B inversion genotypes had east (CCE) foraging phenotypes (Fig. 2D; Fig. S3). Interestingly, we also observed a high frequency of heterozygous individuals (A/B 45% of total) with a roughly even split between the two destination phenotypes (SCS:45%, CCE:55%). Finally, all of the 10 independent validation CCE individuals displayed B/B (50%) or A/B (50%) inversion genotypes (Fig. 2D), aligning with predictions from the original dataset and strengthening support for the role of this putative inversion in driving migratory destination phenotype.

### (d) Genomic variants are associated with neurodevelopment and metabolism

The 111 outlier SNPs associated with migratory destination phenotype were linked to 31 genes, 15 of which have known functional annotations. One of these genes, *Mocos,* is contained entirely within the chromosome 2 putative inversion region, and 67 (60%) of the outlier SNPs fell within or downstream of this gene. This gene encodes Molybdenum cofactor sulfurase, which is involved in regulation of multiple biological processes important to long-distance migration, including metabolism (Table S2). Enrichment analysis did not identify any statistically significant overrepresented pathways, likely due to the small number of 15 genes used as input [56], it identified shared biological processes among these genes. The most common biological processes included synapse organization (GO:0050808), regulation of body fluid level (GO:0050878), sensory organ development (GO:0007423), and membrane trafficking (R-HSA- 199991).

Orange dots represent the independent CCE validation samples. D) Pie charts of each inversion genotype; numbers within pie indicate number of individuals with either migratory destination phenotype possessing each inversion genotype - purple indicates west foragers, blue indicates east foragers. The orange circles beneath the pie charts represent the number of individuals in the independent CCE validation sample set with each inversion genotype.

### (e) Reproductive timing is polygenic

Regions of genomic differentiation associated with reproductive season were more widely distributed across chromosomes and had higher maximum F_st_ values compared to migratory destination phenotype (Fig. 3A; Table S3) . We identified 1120 and 349 outlier SNPs with X^T^X and C_2_ statistics, respectively, with 149 SNPs shared between F_st_, X^T^X, and C_2_ analyses. Absolute nucleotide divergence (D_xy_) was high (99.9^th^ percentile) for 89% of the 149 outlier SNPs (median = 0.62; 0–1 scale; Table S3).

**Figure 3.**
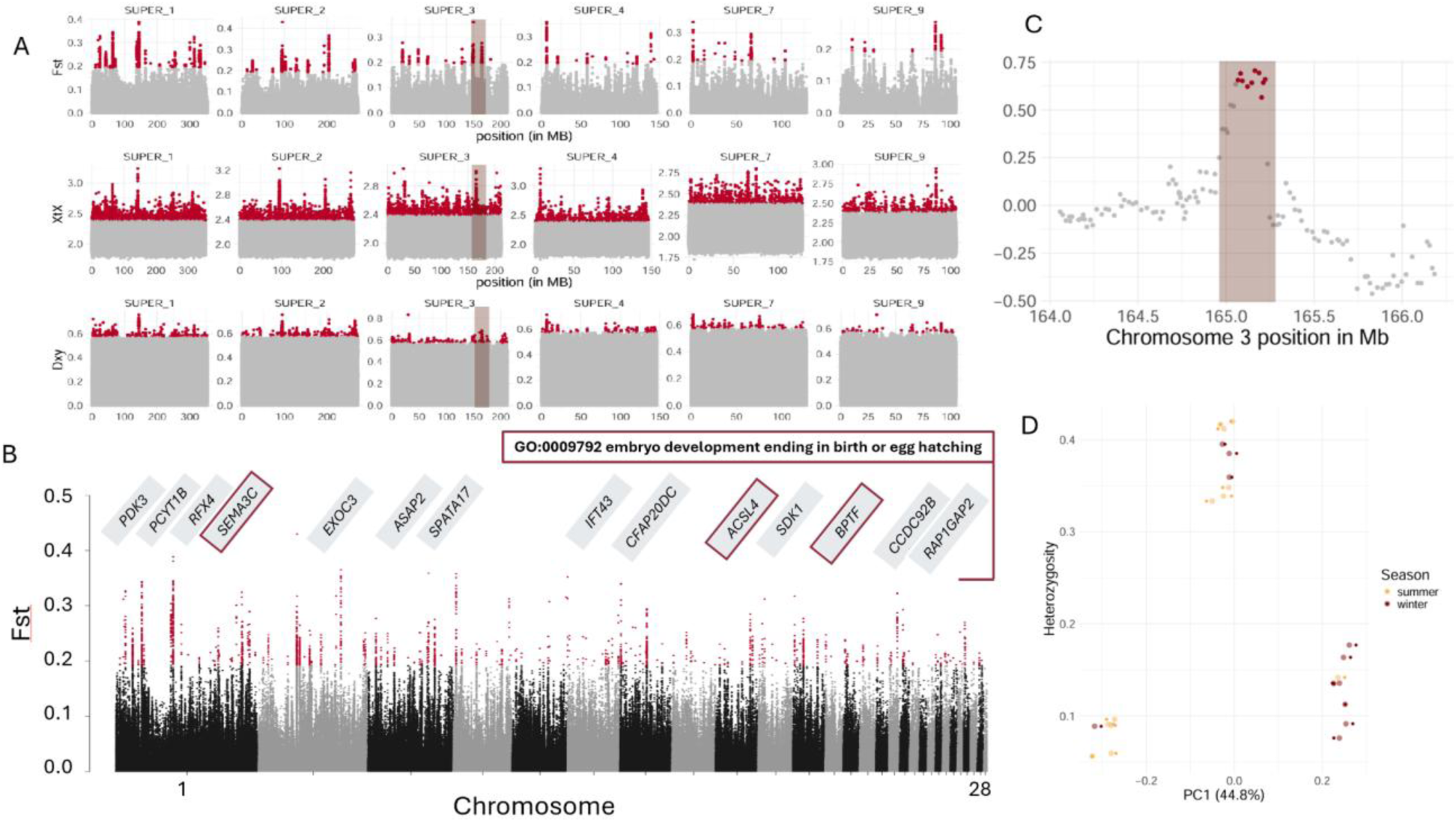
For reproductive timing phenotype A) Triplots for chromosomes with identified genomic islands of divergence. Maroon points indicate outlier regions, light brown box denotes putative inversion region. B) F_st_ across the entire genome, maroon points indicate outliers >= 99.9% of F_st_ values. Genes associated with outlier regions (excluding those in intergenic regions) are shown at the top on their respective chromosomes. Genes outlined in red are associated with the GO biological process for embryo development ending in birth or egg hatchling. C) MDS plot of the putative chromosome 3 (Chr3_2) inversion, x-axis is genome position in Mb. Maroon points indicate 50bp windows where outlier SNPs occur, light brown box indicates putative inversion region. D) PC1 (x-axis) of SNPs within the Chr3_2 putative inversion vs heterozygosity (y-axis) computed over the putative inversion region per individual. Individuals fall into three clusters, with the left cluster representing one homozygous inversion genotype, the middle cluster with high heterozygosity indicating individuals with heterozygous inversion genotype, and the right-most cluster indicating the alternate homozygous inversion genotype.

Of the ten regions with clear GID (two regions on chromosome 1, three regions on chromosome 2, two regions on chromosome 3, single regions on chromosomes 4, 7, and 9), only one region on chromosome 3 appeared to be a putative inversion using our criteria (Fig. 3A,C). This putative inversion of intermediate size spanned 165Kb (0.07% of chromosome 3), exhibited a clear elevated LD block, and displayed a distinct three-cluster pattern of individuals by heterozygosity along PC1 (Fig. 3D; local PCA explained 44.8% (PC1) and 7.0% (PC2) of variance). Other regions exhibited elevated LD but not the expected PCA clustering pattern; eight regions did not exhibit clear elevated LD blocks or expected PCA clustering. Collectively these results suggest that reproductive timing phenotype is driven by polygenic genomic architecture including a single smaller putative inversion along with dispersed regions of genomic differentiation.

### (f) Reproductive timing genomic variants are associated with neural circuitry and reproduction

Outlier SNPs were associated with 65 genes, 36 of which have functional annotations. Shared biological processes included cell-cell adhesion via plasma-membrane adhesion molecules (GO:0098742), protein-containing complex localization (GO:0031503), striated muscle tissue development (GO:0014706), and adenylate cyclase-modulating G protein-coupled receptor signaling pathway (GO:0007188; (Table S3). Embryo development ending in birth or egg hatching (GO:0009792) was the most common shared biological process when using only genes associated with SNPs within introns and exons (Table S3), and the putative inversion on chromosome 3 entirely encompasses the *Spata17* gene (spermatogenesis associated 17), aligned with reproductive biological processes underlying this phenotype. Like the migratory destination phenotype, genes involved in neural circuitry and development processes also play a central role in reproductive timing; 16 of the 36 genes were part of biological processes related to synapse organization and neurodevelopment.

## DISCUSSION

Long-distance oceanic migration is an emblematic life history trait in sea turtles, yet the evolutionary and ecological drivers of these complex behaviors are still largely an enigma. Our study indicates that two key migratory traits —reproductive timing and foraging ground destination—are at least partially driven by genomic determinants in leatherback turtles.

Intriguingly, although both traits display complex genomic architectures involving structural variants and other genomic islands of divergence (GID), migratory destination is largely associated with one intermediate inversion with few additional loci in other regions, while reproductive timing is associated with many outlier regions distributed more throughout the genome. These findings align with recent studies [10,57] supporting that complex eco-behavioral phenotypes can be driven by oligogenic or polygenic mechanisms in natural populations with on- going gene flow. The specific genes associated with migratory phenotypes here have not previously been identified in other migratory taxa, though higher-order biological processes are conserved across leatherbacks and other avian and fish species [9,20,58,59]. Collectively, our results provide the first insight into the genomic mechanisms underlying sea turtle migration, and advance our broader understanding of the evolution of these complex eco-behavioral traits across animal lineages.

The genomic architecture of complex eco-behavioral phenotypes can be highly dependent on population demographic parameters, particularly rates of gene flow. In instances where gene flow is low, locally adapted traits can be maintained by highly polygenic architecture, whereas populations with high gene flow often exhibit fewer, more highly differentiated clusters of loci because beneficial alleles must be protected from recombination to persist [57]. One way this can occur is through the clustering of alleles within structural variants such as genomic inversions [60], which have been increasingly shown to play important roles in maintaining complex traits in natural populations [16,61,62]. While both western Pacific leatherback turtle migratory destination and reproductive timing in this admixed population indeed appear to be controlled by a combination of structural variants and other GID, there are clear differences between the genomic architectures of the traits. Migratory destination in summer nesters is strongly influenced by the putative inversion on chromosome 2 and only a few other loci outside this region, in concordance with expectations for the maintenance of divergent phenotypes with high gene flow. This also aligns with biological expectations, as the individuals foraging in the SCS and CCE are all summer nesters, returning to Indonesian breeding grounds where they would overlap and have opportunity to interbreed. In contrast, reproductive timing appears to be associated with at least nine loci widely distributed across the genome and a smaller putative inversion on chromosome 3; this mixed architecture is more characteristic of adaptive traits maintained under a scenario of intermediate gene flow [57]. Although prior analyses did not detect divergence between summer and winter nesters using genome-wide SNPs, the temporal separation in reproduction may lower gene flow among the two seasonal breeding phenotypes [27]. Interestingly, winter nesters forage in a separate hemisphere from summer nesters [26], likely reducing opportunities for mating between the ecotypes away from the nesting rookeries as well. Thus, it is also possible that reproductive isolation between the phenotypes is occurring, but genetic drift has not had enough time to impart a clear signal of neutral divergence, as sea turtles can exhibit slow mutation rates, long generation times, and demographic history that obscure contemporary demographic isolation [63,64]. Continued population monitoring and genomic analyses of these ecotypes is needed to determine the extent of contemporary gene flow between winter and summer nesting turtles, and more broadly to understand the evolutionary origin and maintenance of the contrasting genomic architectures between these migratory traits.

Complex behavioral phenotypes such as migration involve multiple biological processes [2], involving the modulation of multiple genes and/or those with pleiotropic effects to drive adaptive divergence in natural systems [65]. Nearly all outlier SNPs we identified for both traits occurred in intron or transgenic regions of the genome (Table S2, S3), supporting that their functional effects may impact the splicing or expression of genes rather than changing protein structure.

More study is needed to confirm this, but similar patterns have been observed underlying ecotypes and complex traits in other species [66]. Additionally, genomic inversions and other structural variants can have particularly strong pleiotropic phenotypic effects [67], facilitating the maintenance of multi-trait variation in populations. In our study, the putative inversion on chromosome 2 in the *Mocos* gene may be a prime candidate for pleiotropic effects on multiple traits associated with migratory behavior because it encodes a key enzyme required for xanthine and aldehyde oxidase enzymes –which are expressed in nearly every tissue – to function [68].

These two oxidases impact a swath of traits, including blood pressure maintenance, gut microbiome, antioxidant capacity, inflammatory processes, fat accumulation, neurotransmission, and muscle performance [69–71]. Such biological effects are directly relevant to migratory physiology: leatherback turtles migrating from the temperate CCE back to the Indonesian nesting grounds travel roughly four times farther each way compared to turtles migrating from the tropical SCS, and thereby likely require larger fat deposits, increased blood flow to muscles over a longer duration of time, and mitigation of oxidative stress from exercise, as seen in other migratory species [72]. Indeed, leatherback turtles migrating from temperate foraging grounds have significantly larger curved carapace widths than those from tropical foraging grounds, indicative of increased body mass prior to nesting migration [26]. CCE foragers also have longer curved carapace lengths than SCS foragers [73]; these differences in body size may afford reproductive advantages, as seen in other populations [74], or may be a trade-off required for undertaking longer migrations [75]. More work is needed to confirm these potential relationships and better understand the roles of environmental selection and genetic-environmental interactions in maintaining these divergent phenotypes. Though beyond the scope of our current study, the strong genotype-phenotype associations we observed provide exciting hypotheses for future examination to fully understand the roles of *Mocos* inversions and possible pleiotropy in sea turtle long-distance migration.

Migratory destination and reproductive timing phenotypes in leatherbacks both involve neural processes, which likely modulate the memory, learning, and sensory processing required for long-distance navigation. Many of the genes identified in the present study are associated with behavioral changes in humans and mice (e.g., *Mocos*, *Adgrl3*, *Dscam*, *Adamts18* ; Table S2, S3), or are essential to learning and memory in other migratory species (*Grm8* Table S3), suggesting a possibility of a neuromodulatory effect in turtles. Similar evidence exists in avian species, where allelic differences contribute to morphological differences in the hippocampus and are associated with differing migratory behaviors [59]. However, the timescale and life stage at which the neural circuitry genes function are unclear here and in other taxa; alterations to such genes could affect expression in response to brief environmental cues, program the fundamental development of neural circuits in hatchlings, or affect the plasticity and learning of adults throughout their lives [76,77]. Though the use of geomagnetic cues for long-distance navigation is well documented in sea turtles, birds, and other taxa [78], the functional characterization of magnetoreception in vertebrates largely remains elusive [3]. Some of the genes associated with the destination phenotype (*Adamts18*, *Dscam*) are dominantly expressed in the brain and retinas of mice [79], and may warrant further exploration for a potential role in a retina-based radical-pair magnetoreception process [3,80].

In addition to neural processes, genes underlying the reproductive timing phenotype had functions crucial to embryonic development and fertility (e.g., *Igfbp5, Dazl, Spata17*, and *Cfap20dc*; Table S3). Such genes warrant future study as they may provide insight into why hatching success is substantially lower for summer nests (25.5%) compared to winter nests (47.1%) after accounting for predation and harvest in this population [81]. These genes may also be candidates for comparative studies within and across sea turtles to advance our understanding of why leatherback turtles have lower hatching success rates relative to other species [82].

Environmental factors, such as temperature and moisture associated with differing nesting seasonality, are likely to interact with reproductive timing genes and additionally influence hatching success [83]. Ultimately, the genes underlying both of these migratory traits do not operate in isolation but likely also interact with epigenetic, physiological, and environmental cues to drive leatherback life-history traits.

The western Pacific metapopulation of leatherback turtles is critically endangered, where fisheries bycatch due to entanglement and other threats have reduced abundance by >90% [25,84]. While fisheries bycatch is a threat for all sea turtle species, the pelagic life history and trans-oceanic migrations between nesting and foraging grounds create a particularly high risk of fisheries gear encounters for Pacific leatherback turtles [25]. Understanding the drivers of migratory behaviors, including the extent of their genomic control or flexibility, can inform temporal and spatial forecasts of fisheries interactions under dynamic environmental conditions. Diagnostic assays could also be developed to identify migratory phenotypes of individuals sampled at nesting beaches or in-water and predict which fisheries they are likely to encounter; for example, individuals homozygous for either chromosome 2 inversion could be assigned as western or eastern Pacific foragers. Given evidence that the migratory traits we examined are under at least partial genetic control, other aspects of migratory behavior, such as migratory path, may be as well [20]; future studies combining telemetry and genomics could improve projections of fisheries interactions along different migratory pathways. Additionally, climate change is affecting productivity at foraging grounds and availability of nesting habitat [85,86], such that the mechanisms driving migratory and other eco-behavioral traits will influence exposure risk and resilience. For example, if distant foraging grounds become less productive but migratory behaviors are under strong genetic control, individuals may be canalized to undergo energetically costly trans-oceanic migrations with little reward, potentially reducing reproductive output and altering remigration intervals [87–89]. Conversely, if these behaviors are highly plastic, individuals may switch strategies to compensate. More work is needed to understand these dynamics, but our results suggest these eco-behavioral phenotypes are at least partially heritable which may limit flexibility. Understanding the extent to which migratory traits are under genomic control, plastic, and/or vary between populations has important implications for translocation, genetic rescue, and captive breeding efforts, as there may be genomic incompatibilities [90] or inadvertent introduction of maladaptive variation [91,92]. Finally, conserving leatherback turtles and sea turtles broadly requires preserving morphological, ecological, physiological, and behavioral diversity within and across species to promote resilience in a rapidly changing ocean. Our study provides the first evidence that at least some of this important eco-behavioral diversity in sea turtles is encoded in the genome, providing a foundation for future work to understand and conserve phenotypic variation.

## CONCLUSION

Here we present the first evidence for genomic associations of multiple migratory phenotypic traits in an iconic marine reptile. We identify key genes associated with each trait, finding that high-level biological processes, but not individual genes, related to neurodevelopment and circuitry are conserved across highly diverged taxa, furthering our understanding of the evolution and maintenance of complex eco-behavioral traits. Finally, the complex genomic architecture underlying both migratory destination and reproductive timing in leatherbacks, including putative inversions, adds to a growing body of literature characterizing the maintenance of divergent phenotypes in natural populations under differing scenarios of gene flow. Identifying the functions of putative migratory genes and disentangling the evolutionary mechanisms giving rise to complex migratory behaviors in reptiles are exciting directions for future work with fundamental and applied ecological, evolutionary, and conservation relevance.

## Data accessibility

All original code has been deposited on Github: (https://github.com/jlastoll/leatherback-migration-genomics). Whole genome resequencing data generated in this project is available at NCBI BioProjectID PRJNA1446838; previously published data: NCBI BioProjectID PRJNA 1254794

## Declaration of AI use

AI was not used for any aspect of this study.

## Author Contributions

Conceptualization, J.A. and L.M.K.; methodology and investigation, J.A., D.H., F.P., E.O., J.A.S., T.E., and S.R.B. and L.M.K; writing - original draft, J.A., writing- review & editing, J.A., A.H.A.T, A.P.P., P.H.D., E.O., J.A.S., T.E., S.R.B., L.M.K. ; funding acquisition, L.M.K, P.H.D.; resources, J.A.S, S.R.B, P.H.D.; supervision, L.M.K.

## Acknowledgments

We are extremely grateful to the Abun Leatherback Project, University of Papua, Manokwari, Indonesia, Creausa Hitipeuw (WWF), and Ricardo Tapilatu (UNIPA) for supporting the prior collection of samples that produced the genomic data used in this study. We thank Garrett Lemons, Erin LaCasella and Amy Frey for assistance accessing and linking the phenotypic and sample data in the MMASTR and other NOAA databases used in this study. For their helpful input on study design, analyses and draft revision, we thank past and present members of the Komoroske Lab, particularly Blair Bentley and Katrina Phillips, as well as BAMPHEE, Toni Lyn Morelli, Craig Albertson, Alex Gerson, and Teresa Pegan. Bioinformatic analyses were conducted using the UMass Unity high-performance computing cluster supported by the Massachusetts Green High Performance Computing Center (MGHPCC).

