## Supplemental Figures and Table legends for "A GENOMIC BASIS FOR TRANS-OCEANIC SEA TURTLE MIGRATION"

**SUPPLEMENTARY MATERIALS**

Table S1. Sample metadata, includes which samples were used in the foraging ground vs timing analysis, and sequencing data.

Table S2. Outlier and annotation information for the migratory destination comparison. The first tab includes a guide to the table. The second tab lists all outlier SNPs with F_st_, C_2_, X^T^X, and D_xy_ values, as well as genomic region and nearest gene. The third tab provides annotation information from the Panther gene ontology database and references for select genes of interest from literature searches, and the fourth tab provides annotation information gathered from Metascape.

Table S3. Outlier and annotation information for the reproductive timing comparison. The first tab includes a guide to the table. The second tab lists all outlier SNPs with F_st_, C_2_, X^T^X, and D_xy_ values, as well as genomic region and nearest gene. The third tab provides annotation information from the Panther gene ontology database and references for select genes of interest from literature searches, and the fourth tab provides annotation information gathered from Metascape.

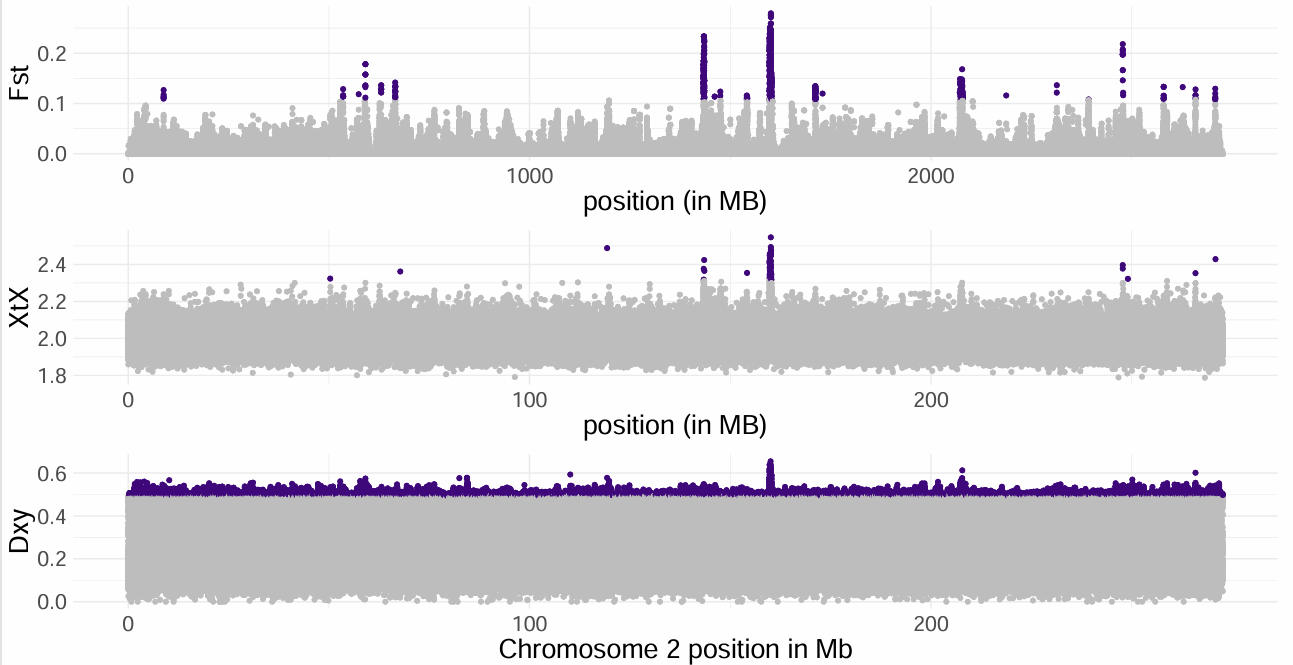

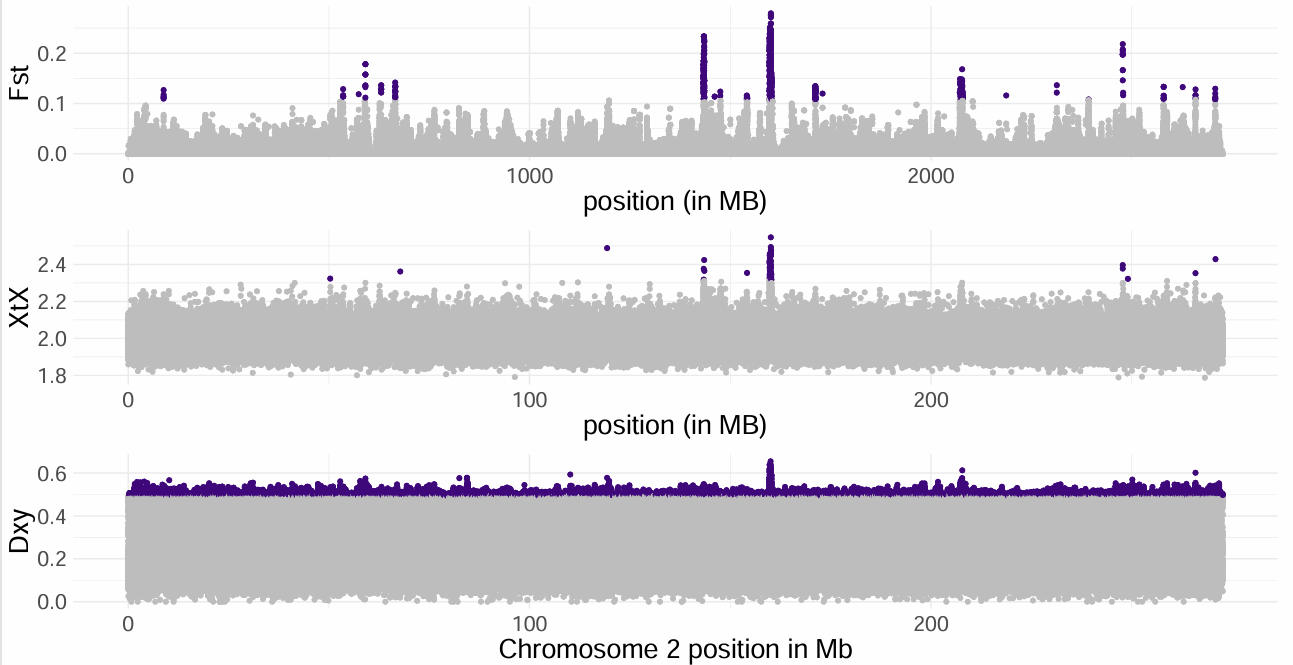

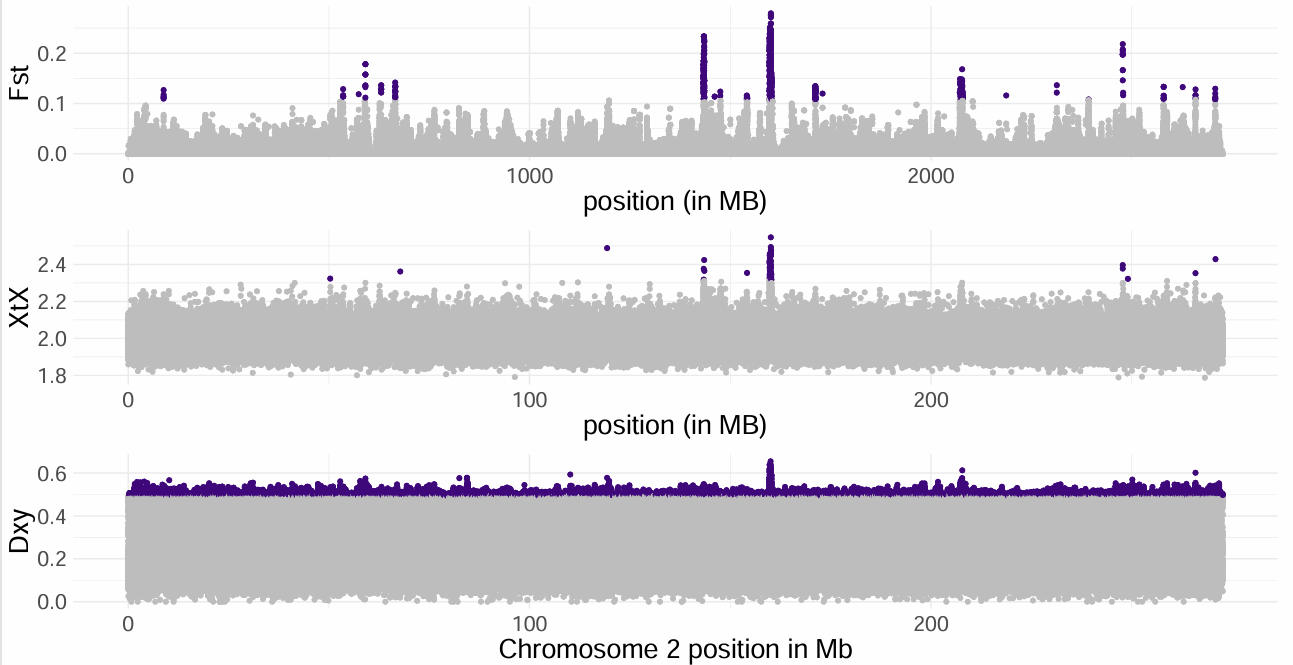

Fig S1. For foraging ground comparison, triplot of F_s_ outliers, X^T^X outliers in middle panel, and D_xy_ per SNP on bottom panel. Purple points indicate outlier regions, which are in the 99.9th percentile for Fst values, or cross thresholds computed based on simulated data for X^T^X and D_xy_ values.

A.

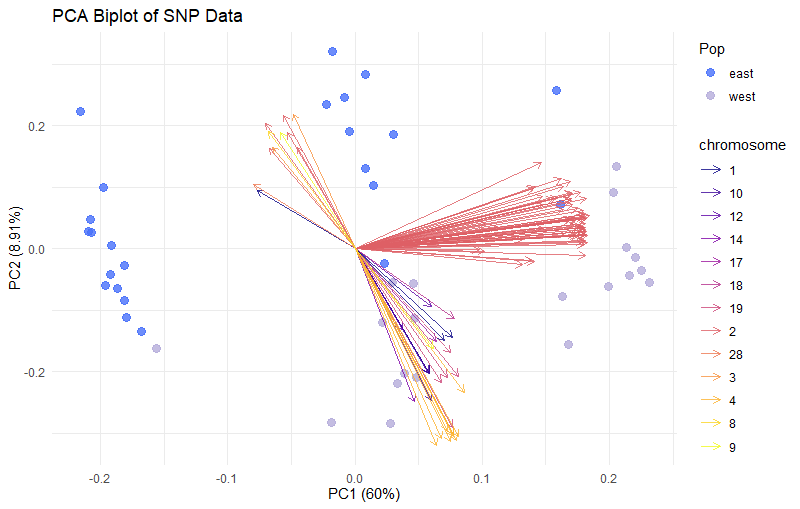

B.

|  | Ref/Ref Genotype Frequency | Ref/Alt Genotype Frequency | Alt/Alt Genotype Frequency |
| --- | --- | --- | --- |
| Eastern Pacific | 0.21 | 0.36 | 0.42 |
| Western Pacific | 0.52 | 0.38 | 0.09 |

Figure S2. We examined allele frequencies at all 111 outlier SNPs to determine if there were any fixed variants for either migratory destination phenotype, or individual SNPs driving phenotype differentiation. A) PCA biplot with loadings of individual SNPs colored by chromosomes. Chromosome 2 SNPs are driving the variation along PC1, no single SNP is driving the divergence in phenotype; B) Mean frequency of each SNP genotype across all 111 outlier SNPs. Homozygous alternate genotypes occurred most frequently in the CCE foraging group and homozygous references genotypes occurred most frequently in the SCS foraging group. We do not observe evidence of a heterozygote disadvantage, though genotypes homozygous for the alternate allele might be selected against in the SCS group.

A.

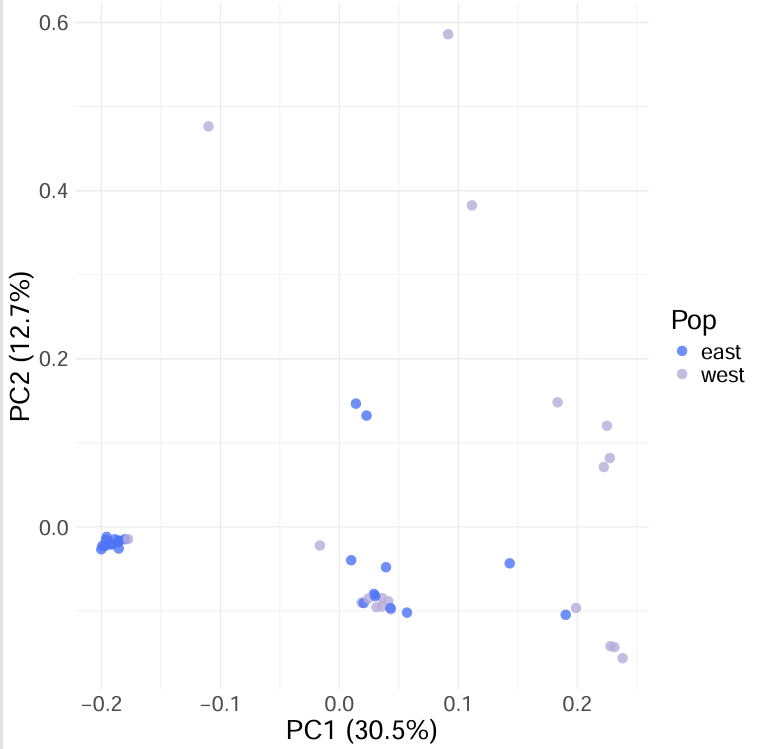

B.

|  | B/B Inversion Frequency | A/B Inversion Frequency | A/A Inversion Frequency |
| --- | --- | --- | --- |
| Eastern Pacific foraging group | 0.54 | 0.38 | 0.08 |
| Western Pacific foraging group | 0.05 | 0.55 | 0.40 |

Figure S3. PCA groupings of SNPs within the putative chromosome 2 inversion. Frequencies of putative chromosome 2 inversion genotypes at hard-called SNPs across foraging groups are given in the table; group sizes are 24 individuals for eastern Pacific foraging group, and 20 individuals for western Pacific foraging group. Note that the single B/B individual in the western pacific foraging group had a lower mapping rate to the reference genome (~80%).

**Supplemental references from Tables S2 and S3:**

Cannarella R et al. 2020 Clinical Evaluation of a Custom Gene Panel as a Tool for Precision Male Infertility Diagnosis by Next-Generation Sequencing. Life 10, 242. (doi:10.3390/life10100242)

Hizawa K, Sasaki T, Arimura N. 2024 A comparative overview of DSCAM and its multifunctional roles in Drosophila and vertebrates. Neuroscience Research 202, 1–7. (doi:10.1016/j.neures.2023.12.005)

Hussan MT, Sakai A, Matsui H. 2022 Glutamatergic pathways in the brains of turtles: A comparative perspective among reptiles, birds, and mammals. Front. Neuroanat. 16. (doi:10.3389/fnana.2022.937504)

Lodjak J, Verhulst S. 2020 Insulin-like growth factor 1 of wild vertebrates in a life-history context. Mol Cell Endocrinol 518, 110978. (doi:10.1016/j.mce.2020.110978)

Miyamoto T et al. 2009 A single nucleotide polymorphism in SPATA17 may be a genetic risk factor for Japanese patients with meiotic arrest. Asian J Androl 11, 623–628. (doi:10.1038/aja.2009.30)

Nie J, Zhang W. 2023 Secreted protease ADAMTS18 in development and disease. Gene 858, 147169. (doi:10.1016/j.gene.2023.147169)

Nongthombam PD, Malini SS. 2023 Association of DAZL polymorphisms and DAZ deletion with male infertility: a systematic review and meta-analysis. Genes Genom 45, 709–722. (doi:10.1007/s13258-022-01345-7)

Rontani P, Perche O, Greetham L, Jullien N, Gepner B, Féron F, Nivet E, Erard-Garcia M. 2021 Impaired expression of the COSMOC/MOCOS gene unit in ASD patient stem cells. Mol Psychiatry 26, 1606–1618. (doi:10.1038/s41380-020-0728-2)

Wang S, DeLeon C, Sun W, Quake SR, Roth BL, Südhof TC. 2024 Alternative splicing of latrophilin-3 controls synapse formation. Nature 626, 128–135. (doi:10.1038/s41586-023-06913-9)
